# Comprehensive integration of single cell data

**DOI:** 10.1101/460147

**Authors:** Tim Stuart, Andrew Butler, Paul Hoffman, Christoph Hafemeister, Efthymia Papalexi, William M. Mauck, Marlon Stoeckius, Peter Smibert, Rahul Satija

## Abstract

Single cell transcriptomics (scRNA-seq) has transformed our ability to discover and annotate cell types and states, but deep biological understanding requires more than a taxonomic listing of clusters. As new methods arise to measure distinct cellular modalities, including high-dimensional immunophenotypes, chromatin accessibility, and spatial positioning, a key analytical challenge is to integrate these datasets into a harmonized atlas that can be used to better understand cellular identity and function. Here, we develop a computational strategy to “anchor” diverse datasets together, enabling us to integrate and compare single cell measurements not only across scRNA-seq technologies, but different modalities as well. After demonstrating substantial improvement over existing methods for data integration, we anchor scRNA-seq experiments with scATAC-seq datasets to explore chromatin differences in closely related interneuron subsets, and project single cell protein measurements onto a human bone marrow atlas to annotate and characterize lymphocyte populations. Lastly, we demonstrate how anchoring can harmonize *in-situ* gene expression and scRNA-seq datasets, allowing for the transcriptome-wide imputation of spatial gene expression patterns, and the identification of spatial relationships between mapped cell types in the visual cortex. Our work presents a strategy for comprehensive integration of single cell data, including the assembly of harmonized references, and the transfer of information across datasets.

**Availability:** Installation instructions, documentation, and tutorials are available at: https://www.satijalab.org/seurat

## Introduction

Recent advances in molecular biology, microfluidics, and computation have transformed the growing field of single cell sequencing beyond routine transcriptomic profiling with single cell RNA-seq (scRNA-seq)^1,2^. Indeed, new approaches now encompass diverse characterization of a single cell’s immunophenotype^3,4^, genome sequence^5,6^, lineage origins^7–9^, DNA methylation landscape^10,11^, chromatin accessibility^12–14^, and even spatial positioning^15–17^. However, each technology has unique strengths and weaknesses, and measures only particular aspects of of cellular identity, motivating the need to leverage information in one dataset to improve the interpretation of another.

The importance of data integration is particularly relevant for approaches that aim to measure distinct modalities within single cells. For example, single cell ATAC-seq (scATAC-seq) can uniquely reveal enhancer regions and regulatory logic, but currently may not achieve the same power for unsupervised cell type discovery as transcriptomics^13,18^. Similarly, methods for multiplexed spatial RNA profiling using *in-situ* hybridization can capture the intricate architecture of tissue organization, but are unable to profile the whole transcriptome^15^. For example, the recently introduced STARmap method enables the measurement of more than 1,000 genes in spatially intact tissue, but forecast this number of genes as an upper limit for such approaches without super-resolution microscopy or the physical expansion of hydrogels^16^. The integration of different single cell technologies with scRNA-seq, such as spatial profiling methods or scATAC-seq, could therefore harmonize these data with transcriptome-wide measurements, allowing not just for the taxonomic listing of cell types, but a deeper understanding of their regulatory logic and spatial organization.

The challenges presented by single cell data integration can be broadly subdivided into two tasks. First, how can disparate single cell datasets, produced across individuals, technologies, and modalities be harmonized into a single reference? Second, once a reference has been constructed, how can its data and meta-data improve the analysis of new experiments? These questions are well-suited to established fields in statistical learning. In particular, domain adaptation aims to identify correspondences across domains to combine datasets into a shared space^19,20^, while transfer learning enables a model trained on a reference dataset to project information onto a query experiment^21,22^. More broadly, these problems are conceptually similar to reference assembly^23^ and mapping^24^ for genomic DNA sequences, and the development of effective tools for single cell datasets could enable similarly transformative advances in our ability to analyze and interpret single cell data.

Recent approaches have established the first steps towards effective data integration. In particular, we recently introduced the use of canonical correlation analysis (CCA)^25^, alongside independent pioneering work leveraging the identification of mutual nearest neighbors (MNNs)^26^, to identify shared subpopulations across datasets. While these approaches can be highly effective, they can also struggle in cases where only a subset of cell types are shared across datasets, or significant technical variation masks shared biological signal. Additionally, these methods focus on scRNA-seq and are not designed to integrate information across different modalities, nor do they enable the transfer of information from one dataset to another.

Here, we present a unified strategy for reference assembly and transfer learning for transcriptomic, epigenomic, proteomic, and spatially-resolved single cell data. Through the identification of cell pairwise correspondences between single cells across datasets, termed “anchors", we can transform datasets into a shared space, even in the presence of extensive technical and/or biological differences. This enables the construction of harmonized atlases at the tissue or organismal scale. These anchors also enable effective transfer of discrete or continuous data from a reference onto a query dataset. This allows for the transfer of cell labels learned from scRNA-seq onto scATAC-seq data to explore differences in the regulatory landscape between distinct interneuron subsets, and the transfer of protein measurements^3^ onto massive public resources to characterize lymphoid populations in human bone marrow. Finally, the anchoring of STARmap and scRNA-seq datasets enables the transcriptome-wide imputation of spatial gene expression patterns in the mouse visual cortex, and the high-resolution spatial mapping of closely related inhibitory and excitatory neuronal subsets. Our results, implemented in an updated version 3 our of open-source R toolkit Seurat, present a framework for the comprehensive integration of single cell data.

## Results

Diverse single cell technologies each measure distinct elements of cellular identity, and are characterized by unique sources of bias, sensitivity, and accuracy^27^. As a result, measurements across datasets may not be directly comparable. For example, expression measurements for scRNA-seq are marred by false negatives (“drop-outs”) due to transcript abundance and protocol-specific biases^27,28^, while expression derived from FISH exhibits probe-specific noise due to sequence specificity and background binding^29^. To address this, we developed an unsupervised strategy to “anchor” datasets together to facilitate integration and comparison. Below we briefly summarize the steps in our approach, alongside a complete description in the Methods, and describe its application to diverse published and newly produced single cell datasets.

### Identifying “anchor” correspondences across single cell datasets

Our motivation for integrating diverse datasets lies in the potential for the information present in one experiment to inform the interpretation of another. In order to relate different experiments to each other, we assume that there are correspondences between datasets, and that at least a subset of cells represent a shared biological state. Inspired by the concept of mutual nearest neighbors, we represent these correspondences as two cells (one from each dataset) that we expect to be defined by a common set of molecular features^26^. While MNNs have previously been identified using L2-normalized gene expression, significant differences across batches can obscure the accurate identification of MNNs, particularly when the batch effect is on a similar scale to the biological differences between cell states. To overcome this, we first jointly reduce the dimensionality of both datasets using diagonalized canonical correlation analysis (CCA), then apply L2-normalization to the canonical correlation vectors (Figure 1A,B). We next search for MNNs in this shared low-dimensional representation. We refer to the resulting cell pairs as “anchors", as they encode cellular relationships across datasets that will form the basis for all subsequent integration analyses. (Figure 1C). Our anchors can successfully recover matching cell states even in the presence of significant dataset differences, as CCA can effectively identify shared biological markers and conserved gene correlation patterns^25^. However, cells in non-overlapping populations should not participate in anchors, representing an important distinction that extends our previous work.

**Figure 1.**
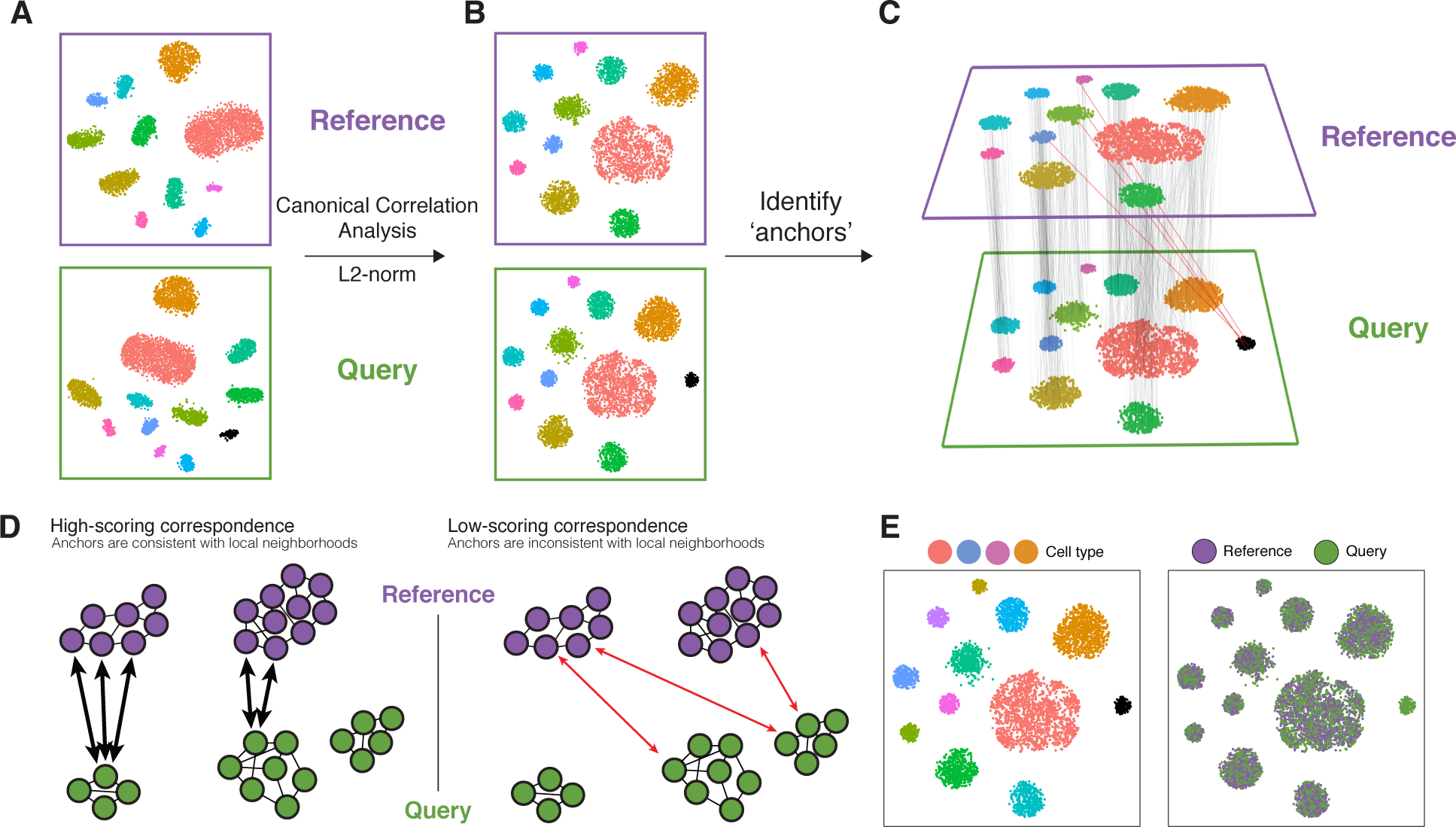
Schematic overview of reference “assembly” integration in Seurat v3. **(A)** Representation of two datasets, reference and query, each of which originates from a separate single cell experiment. The two datasets share cells from similar biological states, but the query dataset contains a unique population (in black). **(B)** We perform canonical correlation analysis, followed by L2-normalization of the canonical correlation vectors, to project the datasets into a subspace defined by shared correlation structure across datasets. **(C)** In the shared space, we identify pairs of mutual nearest neighbors across reference and query cells. These should represent cells in a shared biological state across datasets (grey lines), and serve as “anchors” to guide dataset integration. In principle, cells in unique populations should not participate in anchors, but in practice we observe “incorrect” anchors at low frequency (red lines). **(D)** For each anchor pair, we assign a score based on the consistency of anchors across the neighborhood structure of each dataset. Incorrect anchors tend to have low scores and so are downweighted in future analyses. **(E)** We utilize anchors and their scores to compute “correction” vectors for each query cell, transforming its expression so it can be jointly analyzed as part of an assembled and integrated reference.

Obtaining an accurate set of anchors is paramount to successful integration. Aberrant anchors that form between different biological cell states across datasets are analogous to noisy edges that occur in K-nearest neighbor (KNN) graphs^30^, and can confound downstream analyses. This has motivated the use of shared nearest neighbor (SNN) graphs^31,32^, where the similarity between two cells is assessed by the overlap in their local neighborhoods. As this measure effectively pools neighbor information across many cells, the result is robust to aberrant connections in the neighbor graph. We therefore introduced an analogous procedure for the scoring of anchors, where each anchor pair was assigned a score based on the shared overlap of mutual neighborhoods for the two cells in a pair (Figure 1D; Methods). High-scoring correspondences therefore reflect cases where many similar cells in one dataset are predicted to correspond to the same group of similar cells in a second dataset, reflecting increased robustness in the association between the anchor cells. While we initially identify anchors in low-dimensional space, we also filter out anchors whose correspondence is not supported based on the original untransformed data (Methods). The identification, filtering, and scoring of anchors is the first step for all integration analyses in this manuscript, including reference assembly, classification, and transfer learning.

### Constructing integrated atlases at the scale of organs and organisms

To assemble a reference of single cell datasets in Seurat v3, we aim to identify a non-linear transformation of the underlying data, so that they can be jointly analyzed, in a process conceptually similar to batch correction. We first identify and score anchors between pairs of datasets (referred to as “reference” and “query” datasets) as described above (Figure 1A-D). As introduced by Haghverdi et al.^26^, the difference in expression profiles between the two cells in each anchor represents a batch vector. Therefore, for each cell in the query dataset, we aim to apply a transformation (correction vector) that represents a weighted average across multiple batch vectors. These weights are determined by two components: a cell similarity score, computed individually for each cell in the dataset, and the anchor score, computed once for each anchor. The cell similarity score is defined by the distance between each query cell and its *k* nearest anchors in principal component space (Methods), prioritizing anchors representing a similar biological state. Consequently, cells in the same local neighborhood will share similar correction vectors. The anchor score prioritizes robust anchor correspondences, as described above. By subtracting these weighted correction vectors from the query gene expression matrix, we compute a corrected query expression matrix that can then be combined with the original reference dataset, and used as input for all integrated downstream analyses including dimensionality reduction and clustering. To extend this procedure to multiple datasets, we drew inspiration from “tree alignment” methods for multiple sequence alignment^33^. Here, we first construct a guide tree hierarchy based on the similarity between all pairs of datasets and proceed with recursive pairwise correction up the tree. The similarity score used to construct the hierarchy is computed as the total number of anchors between a pair of datasets, normalized to the total number of cells in the smaller dataset of the pair. This extension for multiple dataset integration was independently conceived but conceptually similar to the Scanorama method^34^.

We hypothesized that our anchoring method could be used to create a reference “atlas” of complex human tissue, by combining diverse datasets across patients, technologies, and laboratories. We therefore examined a collection of eight previously published datasets using tissue from human pancreatic islets, spanning 27 donors, five technologies, and four laboratories^35–39^. Before correcting for technical differences, the cells separated by a combination of dataset of origin and cell type, hindering downstream analysis (Figure S1A). After applying our integration procedure, technical distinctions between datasets were effectively removed (Figure S1B), while major and minor cell populations could be identified through unsupervised graph-based clustering (Figure S1C,D). In addition to reliably detecting all major cell classes present in all datasets (alpha, beta, delta, gamma, acinar, and stellate), we also detected a set of extremely rare cell populations in a subset of the datasets that could not be robustly identified through individual unsupervised clustering analyses (epsilon, schwann, mast and macrophage; Figure S1C).

To examine the robustness of our method to non-overlapping populations, we removed all instances of one cell type from each dataset (e.g. we removed all alpha cells from the celseq dataset, all beta cells from the SMART-seq2 dataset, etc.; Table S1). We then repeated the same integration analysis, and obtained highly concordant results after applying this perturbation (Figure 2A, Figure S2A). Our robustness here originates in part from the anchor scoring approach, as we observed that erroneous anchors in which the query and reference cells belong to different clusters were assigned lower scores compared to consistent anchors and therefore were given less weight in the resulting transformation (Figure 2I).

**Figure 2.**
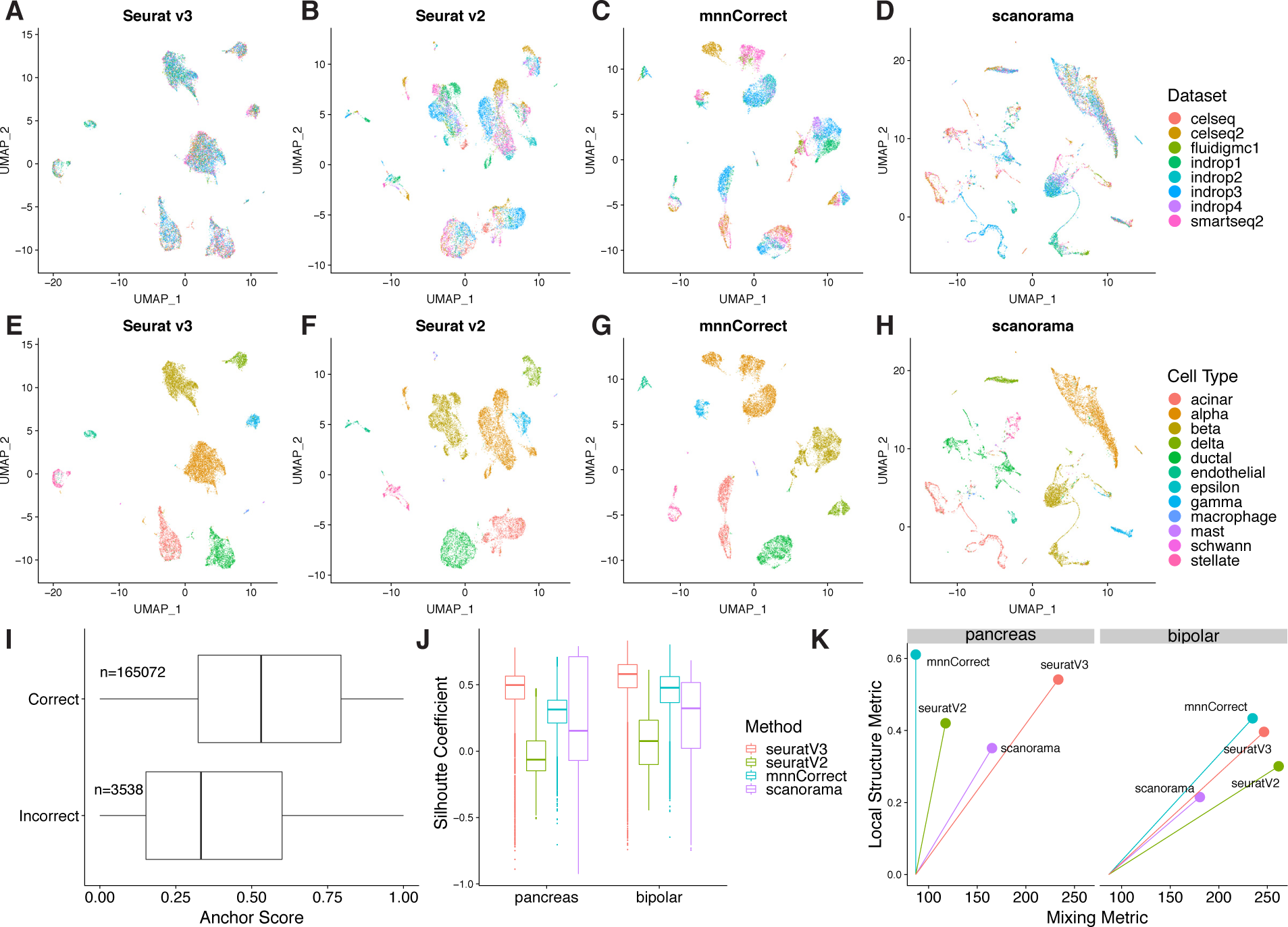
Comparison of multi-dataset integration methods for scRNA-seq. **(A-H)** UMAP plots of eight pancreatic islet cell datasets colored by dataset (A-D) and by cell type (E-H) after integration with Seurat v3, Seurat v2, mnnCorrect, and Scanorama. To challenge the methods’ robustness to non-overlapping populations, a single cell type was withheld from each dataset prior to integration. **(I)** Distribution of anchor scores, separated by incorrect (different cell types in the anchor pair) and correct (same cell type in the anchor pair) anchors. Anchors are from the analysis in Figure S1A. **(J-K)** Metrics for evaluating integration performance across the four methods on two main properties: Silhouette coefficient based on known cell type labels, metrics for cell “mixing” across datasets, and the preservation of within-dataset local structure (Methods).

Using these perturbed datasets, we next benchmarked the performance of our Seurat v3 integration procedure against existing methods (Figure 2A-H). For each tool, we aimed to quantify how well mixed the datasets were after integration, and how well they preserved the structure present in the original datasets (Methods). Methods that perform well in both metrics effectively match populations across datasets without blending distinct populations together. We also calculated silhouette coefficients based on our predefined labels, which captures elements of both sample mixing and local structure. Seurat v3 exhibited the highest silhouette scores and performed well on all other metrics (Figure 2J,K). We obtained equally positive results and benchmarks when examining six batches of murine bipolar cells, which have previously been demonstrated to exhibit batch effects^32^ (Figure 2J,K; Figure S2). We conclude that our anchoring procedure can effectively integrate diverse scRNA-seq datasets and outperforms existing strategies for data integration.

We also considered the potential for our procedure to construct atlases not only at the level of individual tissues, but across entire organisms. To test this, we considered recently published datasets from *Tabula Muris* ^40^, which aimed to profile a diverse set of murine tissues using plate (SMART-Seq2) and droplet (10x Genomics) based assays. These data represent an enormously valuable community resource, but the utility of a single atlas requires that the datasets be harmonized. We identified anchors across 97,029 single cells, representing 18 tissues (12 tissues were represented in both datasets, 6 were only profiled using SMART-seq2), and applied these to integrate the datasets. Integrated visualization revealed extensive mixing of shared cell populations across the two technologies (Figure S3A,B), but cells from the six non-overlapping tissues were not mixed and retained their structure from the original dataset (Figure S3C,D). In particular, we note that this harmonized resource provides exceptional power to detect rare populations, such as tissue-resident plasmacytoid dendritic cells (0.07% cells, detected in 9 tissues), and mesothelial cells (0.05%, detected in 5 tissues), that could not be robustly identified in individual dataset analysis (Figure S3C,E,F). These results suggest an analytical path forward when similar atlas-scale datasets are generated across human tissues with diverse technologies^41^.

### Leveraging anchor correspondences to classify cell states

We next extended our method to transfer information from a reference to a new query dataset. We reasoned that anchors could be used to transfer discrete and continuous data onto query datasets, without requiring modification of the reference. We first considered the problem of cell state classification, where discrete cell labels are learned from pre-trained and reference-derived models, rather than being discovered *de novo* by unsupervised analysis.

As with dataset integration, we approached the classification problem by first identifying anchors between the reference and query datasets. We use the same procedure to identify anchors, with the option to define our search space by projecting a previously computed reference PCA structure onto the query data as opposed to using CCA (Methods). Projecting a query dataset onto an existing PCA structure is more efficient in cases where the query and reference datasets do not exhibit substantial batch differences, when working with a large reference dataset, or when classifying a homogeneous query population. Once we identified anchors, the annotation of each cell in the query set is achieved using a weighted vote classifier based on the reference cell identities, where the weights are determined by the same criteria used in computing the correction vectors for integration (Figure 3A). Since multiple anchors will contribute to the classification of each query cell, these predictions are informed by a cell’s local neighborhood, which increases the overall robustness of the classification call. Additionally, this approach provides a quantitative score for each cell’s predicted label. Cells that are classified with high confidence will receive consistent votes across anchors whereas cells with low confidence, including cells that are not represented in the reference, should receive inconsistent votes and therefore lower scores.

**Figure 3.**
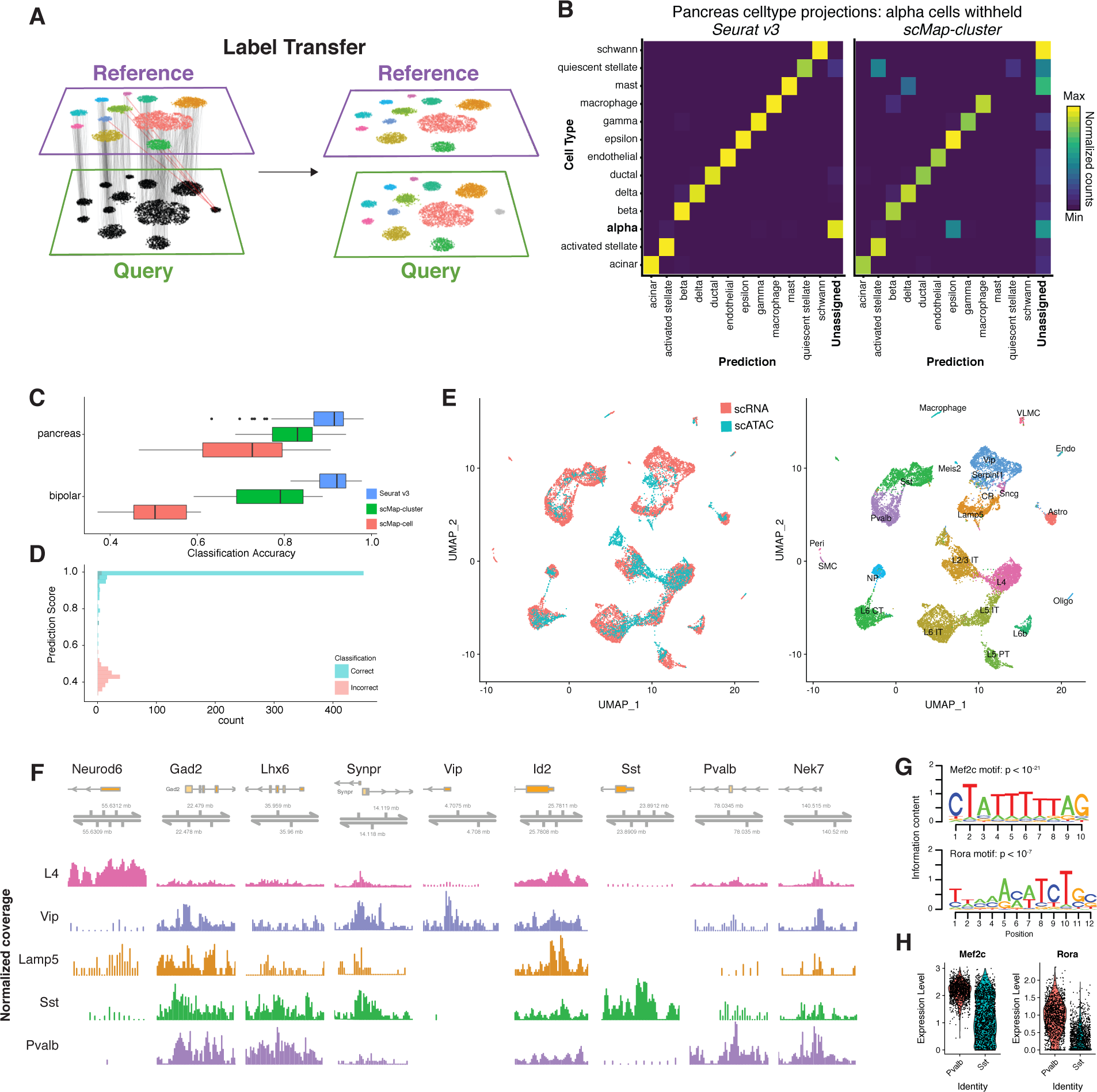
Transferring cell state classifications across datasets. **(A)** Schematic representation where identified anchors allow for the transfer of discrete labels between a reference and query dataset (classification). **(B)** Confusion matrix for one cell type hold-out evaluation where pancreatic alpha cells were removed from the reference. Alpha cells in the query consistently receive the lowest classification score, and are labeled as “Unassigned”. **(C)** Classification benchmarking on 166 test/training datasets from human pancreatic islets and mouse retina. In each evaluation, one cell type was removed from the reference. **(D)** Distribution of prediction scores for one cell type hold-out experiment (as in B). Mis-classification calls are associated with lower prediction scores. **(E)** Joint visualization of scRNA-seq data with classified scATAC-seq cells (left). We identified anchors between scRNA-seq data (reference) and a gene activity matrix derived from scATAC-seq (query) datasets from the mouse visual cortex, and transferred class annotations (right). **(F)** We created pseudo-bulk ATAC-seq profiles by pooling together cells with for each cell type. Each cell type showed enriched accessibility near canonical marker genes. Chromatin accessibility tracks are normalized to sequencing depth (RPKM normalization) in each pooled group. Y-axes for each track were fixed to a minimum of 0 and maximum 200, except for *Id2*, which was fixed to minimum 0 and maximum 400 for each track. **(G)** We searched for overrepresented DNA motifs present in PV-specific accessibility peaks, and identified the Mef2c and Rora motifs as the most highly enriched motifs (p < 10^ࢤ21^ and p < 10^ࢤ7^). **(H)** Both *Mef2c* and *Rora* also exhibit upregulated expression in PV interneurons from scRNA-seq.

We tested our classification in Seurat v3 alongside recently proposed solutions leveraging correlation and nearest-neighbor based classifiers: scMap-cluster and scMap-cell^42^. Using the pancreatic islet and retinal bipolar datasets previously described (Figure 2), we constructed 166 evaluation cases by splitting data into training (reference) and test (query) sets. In each case, we also removed instances of a single cell population (withheld class; for example, alpha cells) from the reference, and then proceeded with classification (Methods). We evaluated classification accuracy by considering the percentage of query cells assigned the correct label, but also examined whether query cells in the withheld class received the lowest classification scores (and were therefore classified as “unassigned"). Seurat consistently received the highest classification accuracy (Figure 3B,C), and correctly assigned low classification scores to query cells that were not represented in the reference (Figure 3B). We note that our increased accuracy stems in part from our ability to use the local neighborhood of a cell (nearby anchors) to increase the robustness of classification, while scMap classifies each cell individually. Additionally, we found that our incorrect predictions were associated with substantially lower classification scores, allowing for the prioritization of high-confidence calls (Figure 3D).

### Projecting cellular states across modalities

We next examined the possibility of applying our classification strategy to transfer cell labels across modalities. For example, we explored whether we could classify individual nuclei from a scATAC-seq dataset based on a reference of transcriptomic states. The potential utility of this approach is underscored by recent studies which have found that scATAC-seq does not currently match the power of scRNA-seq for unsupervised discovery of cell states, including a recently generated scATAC-seq landmark resource of >100,000 nuclei from 13 mouse tissues^18^. For example, 3,482 cells from the prefrontal cortex revealed a cluster of inhibitory interneurons, representing an exciting resource for studying the chromatin accessibility landscape of inhibitory vs. excitatory neurons, but could not identify the well-characterized interneuron subdivisions. Importantly, the authors derive a “gene activity matrix” from the scATAC-seq profiles, utilizing observed reads at gene promoters and enhancers as a prediction of gene activity^43^, representing a synthetic scRNA-seq dataset to leverage for integration.

We reasoned that if we could successfully transfer scRNA-seq derived class labels onto scATAC-seq profiles, we may be able to reveal finer distinctions among the cell types. We therefore considered a deeply sequenced SMART-Seq4 scRNA-seq reference dataset (14,249 cells) of the mouse visual cortex from the Allen Brain Atlas^44,45^, and identified anchors between the scRNA-seq and scATAC-seq using the gene activity matrix derived from scATAC-seq profiles. Joint visualization of the two datasets (Methods) suggested that similar levels of diversity could be identified through integration (Figure 3E). Indeed, by transferring the previously published scRNA-seq celltype labels, we were able to confidently classify 2,548 scATAC-seq cells (projection score > 0.5), into 17 clusters, including eight excitatory and four inhibitory populations (Table S4). Our classifications were consistent with the published labels derived from unsupervised analysis, but revealed substantially increased diversity. For example, 96% of the previously annotated inhibitory neurons were classified as inhibitory in our analysis, but were split into four groups, representing both medial ganglionic eminence (MGE)-derived (SST and PV subsets), and caudal ganglionic eminence (CGE)-derived (Vip and Lamp5) subsets. Pooling nuclei within each projected class together, we obtained pseudo-bulk ATAC-seq profiles. This revealed cell-type specific regulatory loci whose accessibility profiles was fully consistent with expected patterns for all inhibitory cells (Gad2), MGE-derived populations (LHX6), and subset-specific markers (*Pvalb*, *Sst*,*Vip*,*Synpr*,*Id2* ; Figure 3F)^46^. We focused on the PV and SST classes, representing to our knowledge the first efforts to derive and compare genome-wide accessibility landscapes for these closely related interneuron subgroups.

We next performed *de-novo* motif analysis in an attempt to discover cis-regulatory DNA sequences that differentially regulate PV and SST interneurons. While few validated regulators that drive specific interneuron fate decisions are known, we have previously shown that the transcription factor *Mef2c* is upregulated in embryonic precursors of PV interneurons, and is specifically required for their development^46^. Strikingly, our scATAC-seq analysis revealed a strong enrichment for Mef-family motifs (including *Mef2c*) in peaks with increased PV-accessibility, representing the highest-scoring motif (Figure 3G). We observed other motifs for putative regulators, including the putative regulator *Rora* ^47^ (second highest motif; Figure 3G). Intriguingly, as with *Mef2c*,*Rora* also exhibits RNA upregulation in PV compared to SST interneurons (Figure 3H), and may also play roles in fate specification. Taken together, these results highlight the role of *Mef2c* and other transcription factors in establishing or maintaining the chromatin landscape necessary to express the functional receptors and transporters that establish the specific identity of PV cells.

Our results demonstrate the potential for transferring scRNA-seq derived annotations onto chromatin accessibility data. We emphasize that this strategy requires an initial step where ATAC-seq data is converted to a predicted gene expression matrix^43^. Existing strategies for this task likely assume that chromatin accessibility is positively correlated with gene expression. While this assumption has generally held true and enabled the prior interpretation of scATAC-seq data in the developed brain^13,18^, there may be cases where accessibility is a poor proxy for transcriptional output, particularly in developing systems where chromatin changes precede gene expression^48^. In these cases, we expect that we would not be able to form consistent anchors across datasets. However, effective integration can occur even if only a subset of features exhibit coordinated behavior across RNA and chromatin modalities, similar to how cross-species scRNA-seq datasets can be effectively integrated even when only a subset of gene expression markers are conserved^25^.

### Transferring continuous and multimodal data across experiments

Though we previously demonstrated how anchors could be utilized to transfer discrete classifications across datasets, we reasoned that the same methods could be used to transfer continuous data as well. This is of particular interest for the growing suite of multimodal single cell technologies that measure multiple aspects of cellular identity. Transfer learning could therefore be used to fill in missing modalites in key datasets. For example, the Human Cell Atlas recently released a freely available resource of 274,599 healthy bone marrow cells from 8 individual donors^49^. This represents an extraordinary community resource to study the human immune system, but does not contain cell surface protein measurements, which could substantially improve the ability to interpret and annotate this resource. We hypothesized that by generating a human bone marrow dataset with our recently developed CITE-seq technology^3^, where immunophenotypes are measured in parallel with transcriptomes using DNA-barcoded antibodies, we could effectively transfer protein expression data to the HCA dataset. Additionally, we highlight that this method can be successful even in the absence of correlation between RNA and protein for individual genes (e.g. between *Cd4* transcript and CD4 protein), though it does require that a combination of genes exhibit expression patterns that are correlated with cellular immunophenotype (e.g, modules of markers for CD4+ T cells).

### Predicting protein expression in human bone marrow cells

We performed a CITE-seq experiment on human bone marrow cells^3^, capturing 33,454 cells for which we measured cellular transcriptomes alongside 25 cell-surface proteins representing well-characterized markers (median 4,575 RNA UMIs and 2,312 ADT UMIs per cell; Table S3, S4; Figure S4). We first performed cross-validation within the CITE-seq data by randomly assigning cells to a reference or query dataset, and identified anchors between them. As with our discrete classifications, we predicted protein levels in the query dataset using a weighted average of CITE-seq counts from the reference anchor cells, which we then compared with the original measurements (Figure 4A).

**Figure 4.**
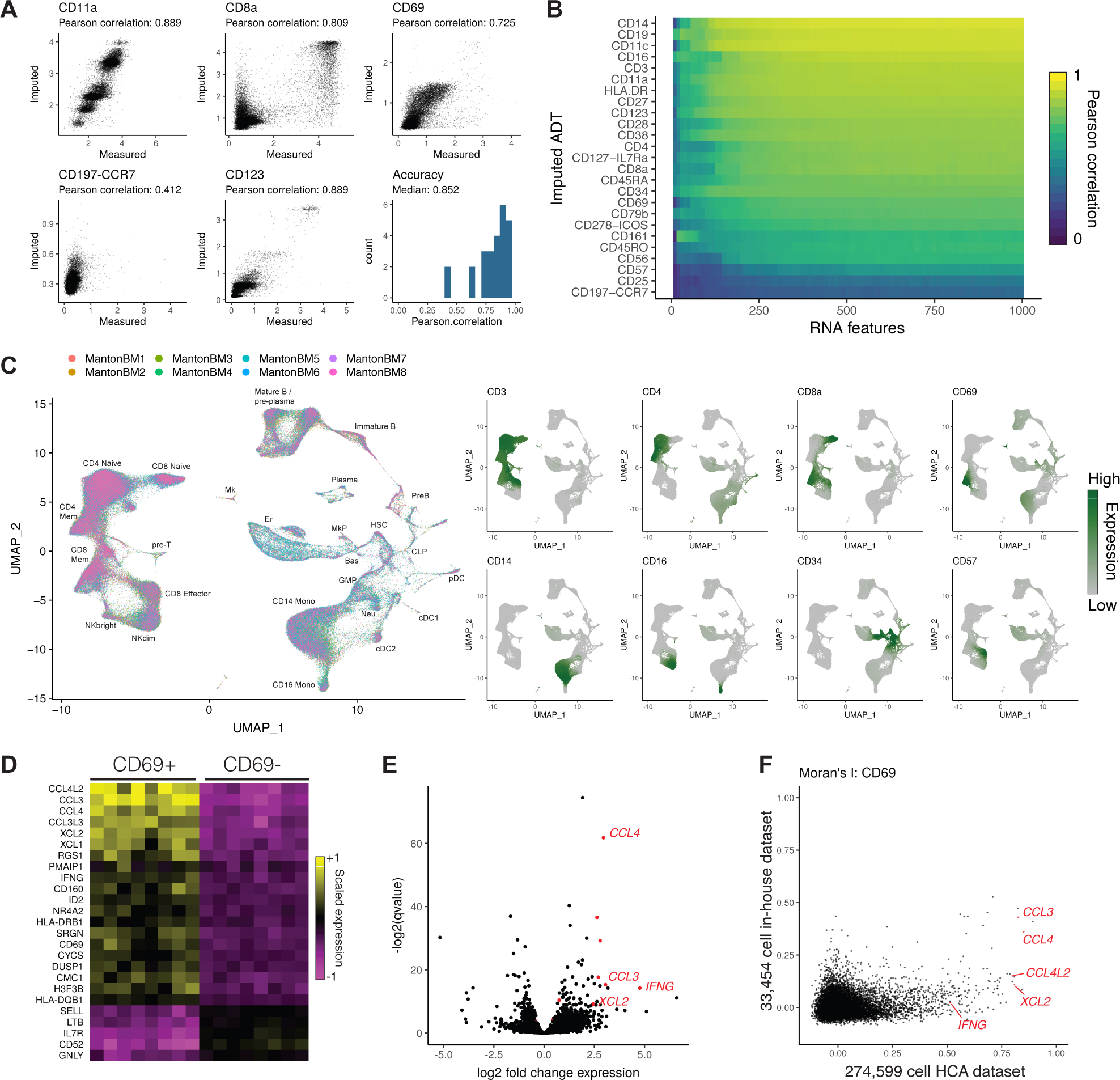
Imputing immunophenotypes in a transcriptomic atlas of the human bone marrow. **(A)** Cross-validations for immunophenotype imputation, performed using a CITE-seq dataset of 35,543 bone marrow cells and 25 surface proteins. **(B)** Prediction accuracy as a function of the number of transcriptomic features used to determine anchors. We progressively downsampled the number of RNA features used to make protein expression predictions, and repeated the cross-validation analysis with each downsampling. **(C)** We integrated 274,599 bone marrow cells produced by the Human Cell Atlas and annotated the integrated cell types. Using the CITE-seq bone marrow cells, we predicted protein expression levels in the integrated HCA dataset, and observed expression patterns consistent with the known cell types. **(D)** Predicted CD8+ CD69+ cells up-regulate a module of inflammatory cytokines and chemokines across all eight donors. Shown are averaged RNA expression values for each human donor. **(E)** We validated CD69+ marker genes identified in the scRNA-seq data by performing bulk RNA-seq on FACS-isolated CD8+ CD69 +/− cells, which revealed a similar set of deferentially expressed genes. **(F)** We ordered CD8+ memory cells by their CD69 expression in the HCA and CITE-seq datasets, and computed the autocorrelation for each gene along this CD69 axis (Moran’s I). CD69+ marker genes consistently showed a higher Moran’s I value in the HCA dataset, reflecting the increased statistical power accompanying an order-of-magnitude greater cell number.

For most proteins (22/25), we observed strong correlation between the measured and imputed expression levels (Figure 4A,B; median R=0.852; Figure S5), with the remaining residual encompassing background CITE-seq binding (perhaps driven by differences in cell size), stochastic variation in protein expression, or technical noise. In the three cases where we observed poor correlations, either poor antibody specificity or a lack of transcriptomic markers that correlate with immunophenotype could explain these results. By downsampling RNA features used to identify anchors and repeating the cross-validations, we found that prediction accuracy began to saturate at approximately 250-500 features (Figure 4B), suggesting that only a subset of shared genes need to be measured across experiments in order to transfer additional modalities across datasets.

Having demonstrated our ability to accurately impute immunophenotypes, we next transferred protein expression data from our CITE-seq experiment to the Human Cell Atlas bone marrow resource of 274,599 cells across eight human donors^49^, after first integrating the eight donor datasets using Seurat v3 to mitigate batch effects among the donors (Figure 4C; Figure S6, S7). Encouragingly, our imputed immunophenotypes were consistent with the well-studied expression patterns of key markers in the hematopoietic system (Figure S8), including high predicted CD34 expression in early hematopoietic progenitors, mutual exclusivity between CD8a and CD4 expression, and canonical marker expression in monocytes (CD14), NK (CD16 / CD56), and B cell (CD19) populations. Intriguingly, we identified a sub-population of CD8+ memory cells marked by sharply elevated predicted expression of CD69 (Figure 4C). While CD69 has been proposed as an early activation marker of T cells^50^, the molecular phenotype and significance of CD8+ CD69+ cells in the bone marrow is not well understood. Recent evidence in particular suggests that CD8+ cells upregulate this marker without accompanying changes in the transcriptome, and that the transcriptome of these cells is in a resting state^51^. We therefore sought to identify genes whose measured expression in the HCA data was associated with predicted CD69 expression.

We observed a clear module of genes associated with increased CD69 expression across all eight human donors (Figure 4D), including cytokines, chemokines, and granzyme molecules, with ontology analysis revealing striking enrichment for genes involved in IFN-*γ* responses (P < 10^−11^; Figure S9). We validated this finding by sorting CD8+/CD69+ and CD8+/CD69-T cells, performing bulk RNA-seq (four replicates each), and observing differential expression of our top markers (Figure 4E). Importantly, while we observed similar CD69+ heterogeneity in an independent analysis of the original CITE-seq dataset (Figure S4), this dataset contained an order-of-magnitude fewer cells, and as a result exhibited substantially lower power to detect genes associated with CD69 expression. To quantify this, we ordered cells in the CITE-seq and HCA datasets along an axis of CD69 expression, and computed Moran’s I statistic, a measure of spatial autocorrelation, for each gene. We consistently observed substantially higher Moran’s I values in the HCA dataset, and could not identify key inflammatory genes (including *IFNG*) as outliers from the CITE-seq data alone (Figure 4F). Further experiments are needed to reveal the functional importance of this population, but notably, secretion of inflammatory cytokines like IFN-*γ* can alter the bone marrow microenvironment and hematopoietic output^52^. Taken together, these results demonstrate how transfer learning can be used to facilitate biological discovery across datasets, and to impute missing modalities in key resources.

### Spatial mapping of single cell sequencing data in the mouse cortex

As a final demonstration of transfer learning using our Seurat v3 method, we explored the integration of multiplexed *in-situ* single cell gene expression measurements (FISH) with scRNA-seq of dissociated tissue. While we and others^53–56^ have previously demonstrated analytical strategies to map single cells to their original spatial position, these strategies require the tissue to have a stereotypical structure, and rely heavily on transcriptional gradients to facilitate the spatial mapping of cells. In principle, the harmonization of multiplexed FISH or *in-situ* RNA sequencing with scRNA-seq would enable similar goals to be achieved for any biological system, a challenge that is of paramount importance to understand the spatial organization and regulation of cells and tissues. While imaging datasets have an upper limit on the number of dimensions that can be simultaneously profiled per cell, our previous results indicated that only a subset of transcriptomic features were necessary to facilitate integration (Figure 4B). We therefore considered two complementary datasets of the mouse visual cortex, the deeply sequenced SMART-Seq4 dataset from the Allen Brain Institute^45^ (as in Figure 3E; 14,249 cells, 45,768 transcripts), and the recently published STARmap *in-situ* gene expression datasets of the same tissue (1,539 and 890 cells, 1,020 genes)^16^.

After identifying anchors between the datasets, we imputed spatial expression patterns across the transcriptome by transferring the expression of all measured scRNA-seq transcripts (45,768) onto the STARmap datasets (Figure 5A). For genes with well-established spatial patterns of expression (for example, the layer-specific marker genes *Lamp5* and *Cux2*), our imputed patterns were fully concordant with the measured STARmap data (Figure 5B, S10A). Similarly, genes that were cell-type specific but not spatially restricted (for example, the interneuron subtype marker *Sst* and endothelial marker *Bsg*) also exhibited identical patterns in the imputed and measured data. However, we also observed cases where the original STARmap data exhibited a weak signal that was strengthened in the imputed data (for example, *Rorb* and *Syt6*). These cases could reflect stochastic cellular expression, technical noise in the STARMap data, or imputation errors - although our predictions here were further supported by an independently derived, highly sensitive cyclic single molecule FISH (osmFISH) experiment^17^ (Figure 5C). By transferring the remaining scRNA-seq genes onto spatially-resolved cells, we were further able to predict spatial patterns for genes that were not originally profiled profiled by STARmap. We identified four representative cases (Figure 5D), each of which contains strong external support in the published literature^57,58^, or the Allen Brain Atlas^44,45^ (Figure S11). Moreover, when repeating the imputation procedure on a second independent STARmap replicate (890 cells), we found that our gene-level predictions for spatial association were highly reproducible, with the exception of a small group of endothelial markers reflecting a non-random spatial distribution (a “strip") of endothelial cells in only the first replicate (Figure 5E; Figure S10A,B).

**Figure 5.**
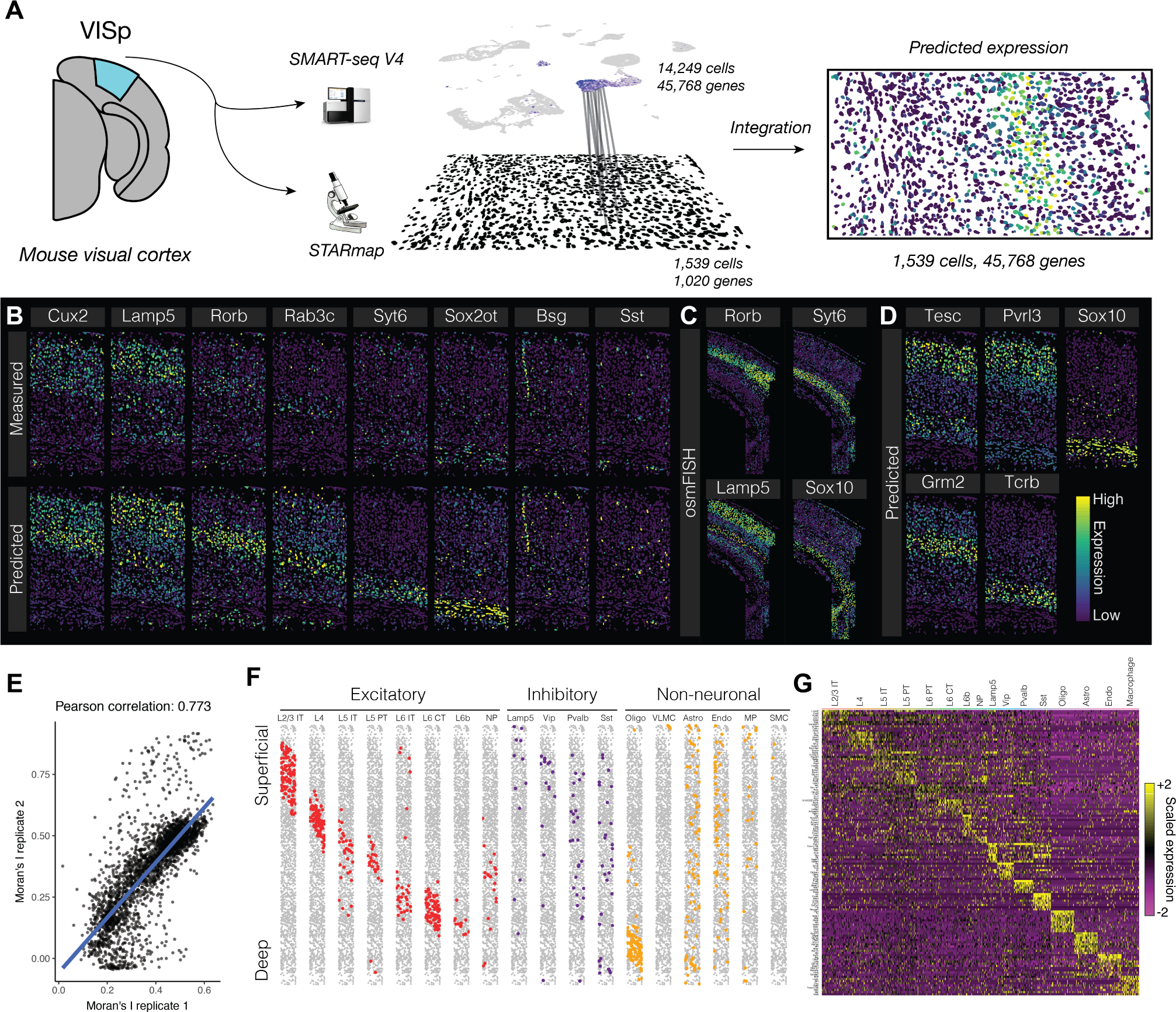
Spatial patterns of gene expression in the mouse brain. **(A)** Schematic representation of data transfer between scRNA-seq and STARmap datasets. After identifying anchors using the subset of genes measured in both experiments, we subsequently transfer sequencing data to the STARmap cells, predicting new spatial expression patterns. **(B)** Leave-one-out cross validation for 8 genes, exhibiting predicted expression patterns, and original STARmap measurements. **(C)** Gene expression patterns for *Rorb*, *Syt6*, *Lamp5* and *Sox10*, as measured by osmFISH, a highly sensitive single molecular assay^17^, in the mouse somatosensory cortex. **(D)** Predicted expression patterns for four genes not originally profiled by STARmap, with external validation in Figure S11. **(E)** Correlation between Moran’s I value, a measure of spatial autocorrelation, for each predicted gene expression pattern in two STARmap replicates. **(F)** Horizontally-compressed STARmap cells with predicted cell type transferred from the SMART-seq4 dataset. **(G)** Expression of cell type marker genes in each predicted STARmap cell type (both replicates combined).

As we previously demonstrated using scATAC-seq data, our anchoring procedure allows us to classify cells across modalities based on scRNA-seq annotations. We therefore transferred cell type labels from the SMART-seq4 dataset to the STARMap cells, classifying 1,915 (79%) of cells with prediction score > 0.5, but conservatively chose to consider the 1,210 (50%) cells with the highest prediction scores for downstream analysis (Figure 5F; Methods). These classifications revealed subdivisions that could not be identified even through iterative clustering of the original dataset. For inhibitory cells, we identified cells from the four major classes (Sst, Pvalb, Lamp5/Id2, Vip), each expressing canonical markers in the original STARmap dataset (Figure 5G). In excitatory cells, we annotated cells from 8 different clusters, representing not only layer-specific populations, but also separating intratelencephalic (IT), pyramidal tract (PT), corticothalamic (CT), and L6b sublayer populations within individual layers (Figure 5F,G).

Lastly, we examined the spatial distribution of our annotated cell-types, searching for non-random patterns. As previously reported, MGE-derived interneurons were enriched in Layers 4/5, CGE-derived interneurons were enriched in layers 1-3, and excitatory populations were strongly associated with individual layers (Figure 5F)^59^. However, after closely examining the mapping patterns, we observed differences in the laminar distributions for neurons even within the same layer, including IT and PT neurons (Layer 5), and IT and CT neurons (Layer 6), suggesting a complex interplay between excitatory specification and within-layer spatial positioning. These results were reproduced in the second STARmap replicate dataset (Figure S10C), but the functional consequences remain to be explored. We conclude that anchoring imaging and sequencing datasets enables the transcriptome-wide prediction of spatial expression patterns, and the harmonization of scRNA-seq derived cell type labels with *in-situ* gene expression datasets. As multiplexed image-based single cell methods and datasets continue to grow and develop, the integration of sequencing and imaging datasets therefore represents a powerful and exciting opportunity to construct high-resolution spatial maps of any biological system.

## Discussion

We have developed a strategy for the comprehensive integration of single cell data, and apply this to derive biological insights jointly from transcriptomic, epigenomic, proteomic, and spatially-resolved single cell data. Our strategy tackles several technical challenges, starting with the unsupervised identification of cell pairs across datasets, deemed “anchors", that represent a similar biological state. This enables us to either assemble multiple datasets into an integrated reference, or to transfer data and metadata from one experiment to another. We anticipate that as single cell RNA-seq experiments have only recently become routine, the challenge of reference assembly will be of particular importance to both small labs and large consortia, as new experiments will continually uncover increasingly rare and subtle biological states. However, as these references begin to stabilize, projecting both discrete labels and continuous data onto new datasets will be of transformative value to the interpretation of new datasets, analagous to how short-read mapping enabled the rise of multiple functional genomics technologies^24,60^. Throughout multiple examples in this manuscript, we demonstrate how integrated analysis can reveal biological insights that require the cluster identification and annotation inherent to scRNA-seq analysis, but could not be identified by any single experiment. In particular, we derive *in-silico* bulk ATAC-seq profiles for finely resolved interneuron subsets whose identities can be classified with the assistance of transcriptomic data, as well as identifying cell surface proteins that can successfully enrich for transcriptomically-defined T cell subsets in human bone marrow. Lastly, we demonstrate how scRNA-seq and *in-situ* gene expression data can be integrated to robustly predict spatial expression patterns transcriptome-wide, and even to identify high-resolution spatial relationships between closely related neuronal subtypes.

We expect our strategy to be broadly applicable to integrate and transfer a broad spectrum of single cell data and phenotypes across experiments. These include additional epigenomic^10–14^, chromosome conformation^61,62^, and RNA modification^63^measurements that are increasingly being profiled at the single cell level, and even computationally derived phenotypes such as RNA velocity^64^. We believe that the integration of sequencing and imaging datasets represents a particularly promising application in the near future. Recent work based on the spatial analysis of protein panels^65,66^ has poignantly demonstrated how changes in tissue organization can dramatically shift across disease states. By integrating single cell transcriptomics with spatial datasets, these analyses can consider not only broadly defined cell types, but subtle alterations in cell state, even for genes that are not directly measured in an imaging probe set. Future extensions could utilize these molecular data to assist in the image alignment of multiple datasets, or even integrate with perturbation screens to help infer causal relationships^67^.

Lastly, our results suggest that single cell RNA-seq can serve as a general mediator for single cell data integration. Not only is its application commercialized and routinely available, but transcriptome-wide gene expression data encodes multiple aspects of cellular identity and “metadata", even if they are lost during the experimental process. Moreover, its intermediate position in the central dogma allows for proximity to multiple molecular processes, including both transcriptional, posttranscriptional, and translational regulation. We therefore suggest that scRNA-seq may serve as a ’universal adapter plug’ for single cell analysis, facilitating integration across multiple technologies and modalities, and enable a deeper understanding of cellular state, interactions, and behavior.

## Methods

### Seurat integration method

The Seurat v3 anchoring procedure is designed to integrate diverse single cell datasets across technologies and modalities. To facilitate the assembly of datasets into an integrated reference, Seurat returns a corrected data matrix for all datasets, enabling them to be analyzed jointly in a single workflow. To transfer information from a reference to query dataset, Seurat does not modify the underlying expression data, but instead projects either discrete labels or continuous data across experiments. While the use-cases for each approach will depend on the user and particular experiment, the underlying methods are conserved across approaches. When possible in the methods, we specify the function in Seurat where the method is implemented, to facilitate users exploring the source code, which is freely available at https://satijalab.org/seurat.

Our approach consists of four broad steps, as explained in detail below: (1) data preprocessing and feature selection, (2) dimension reduction and identification of “anchor” correspondences between datasets, (3) filtering, scoring, and weighting of anchor correspondences, (4) data matrix correction, or data transfer across experiments.

### Parameters for Seurat v3 integration

To exemplify the general utility of our approaches, we aimed to minimize the free parameters that can be tuned for each analysis and to utilize default parameters in all cases. All parameters are described throughout the methods, even when their default values are fixed for all analyses in this manuscript.

One parameter we expect to fluctuate across across datasets represents the estimated “dimensionality” of the data. This affects, for example, the number of principal components or canonical correlation vectors that are calculated during dimensional reduction. Larger datasets will typically have increased dimensionality, particularly if they represent increasingly heterogeneous populations. While we have previously suggested using saturation or statistical-resampling based approaches to estimate dataset dimensionality^25^, a robust fully unsupervised procedure to identify this value remains a fundamental challenge in the analysis of high-dimensional data. Here, we neglect to finely tune this parameter for each dataset, but still observe robust performance over diverse use cases. For all neuronal, bipolar, and pancreatic analyses we choose a dimensionality of 30. For scATAC-seq analyses we chose a dimensionality of 20. For analyses of human bone marrow and the integration of mouse cell atlases, we choose a dimensionality of 50 and 100 respectively, representing the significant increase in dataset size and heterogeneity for these cases.

We also allow for the use of approximate nearest neighbor methods, using the RANN package in R^68,69^. While not enabled by default, the user can set the error bound parameter (nn.eps) to increase the speed of nearest neighbor identification. This parameter is set to 0 by default, but for analyses where more than 50,000 cells are analyzed in total (Figure 4, Figure S3), we set this value to 1. Unless otherwise specified, all other quantitative parameters are fixed to default values.

### Data preprocessing

#### Normalization

For all analyses, we employed standard pre-processing for all single cell RNA-seq datasets. Unless otherwise specified, we first performed log-normalization of all datasets, using a size factor of 10,000 molecules for each cell. We next standardized expression values for each gene across all cells (z-score transformation), as is standard prior to running dimensional reduction tools such as principal component analysis. These steps are implemented in the NormalizeData and ScaleData functions in Seurat.

#### Feature selection for individual datasets

In each dataset, we next aimed to identify a subset of features (i.e. genes) exhibiting high variability across cells, and therefore represent heterogeneous features to prioritize for downstream analysis. Choosing genes solely based on their log-normalized single cell variance fails to account for the mean-variance relationship that is inherent to single cell RNA-seq. Therefore, we first applied a variance-stabilizing transformation to correct for this, as first outlined by Mayer, Hafemeister & Bandler et al.^46^.

To learn the mean-variance relationship from the data, we computed the mean and variance of each gene using the unnormalized data (i.e. UMI or counts matrix), and applied log_10_-transformation to both. We then fit a curve to predict the variance of each gene as a function of its mean, by calculating a local fitting of polynomials of degree 2 (R function loess, span = 0.3). This global fit provided us with a regularized estimator of variance given the mean of a feature. As such, we could use it to standardize feature counts without removing higher-than-expected variation.

Given the expected variances, we performed the transformation

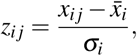

where *z*_*ij*_ is the standardized value of feature *i* in cell *j*, *x*_*ij*_ is the raw value of feature *i* in cell *j*, 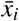 is the mean raw value for feature *i*, and σ_*i*_ is the expected standard deviation of feature *i* derived from the global mean-variance fit. To reduce the impact of technical outliers, we clipped the standardized values to a maximum value of 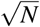, where *N* is the total number of cells. For each gene, we then computed the variance of standardized values across all cells. This variance represents a measure of single cell dispersion after controlling for mean expression, and we use it directly to rank the features. Unless otherwise noted, we selected the 2,000 genes with the highest standardized variance as “highly variable". This procedure is implemented in the FindVariableFeatures function in Seurat v3 (method=”vst”).

#### Feature selection for integrated analysis of multiple datasets

When performing integration across datasets, we aimed to give priority to features that were identified as highly variable in multiple experiments. Therefore, we first performed feature selection on each dataset individually, using the procedure described above. We next prioritized features across multiple experiments by examining the number of datasets in which they were independently identified as highly variable. From this ranked list of features, we took the top 2,000 to use as input for downstream analyses. We broke ties by examining the ranks of the tied features in each original dataset and taking those with the highest median rank. These steps are implemented in the SelectIntegrationFeatures function in Seurat v3.

### Identification of anchor correspondences between two datasets

A key step for all integration analyses in this manuscript is the unsupervised identification of “anchors” between pairs of datasets. These anchors represent two cells (with one cell from each dataset), that we predict to originate from a common biological state. Anchors for reference assembly or transfer learning are calculated using the FindIntegrationAnchors and FindTransferAnchors functions, respectively, in Seurat v3.

We initiate this process through dimensional reduction, aiming to place datasets in a shared low-dimensional space. For reference assembly, we utilize canonical correlation analysis (CCA) as an initial dimensional reduction. As we have previously demonstrated^25^, the canonical correlation vectors described by CCA effectively capture correlated gene modules that are present in both datasets, representing genes that define a shared biological state. In contrast, principle component analysis (PCA) will identify sources of variation even if they are only present in an individual experiment, particularly if there are significant technical effects across experiments. We therefore utilize CCA when integrating scRNA-seq datasets into a common reference, or when identifying anchors from single cell data spanning modalities.

Canonical correlation vectors are calculated as described previously^25^. Briefly, let *X*_*f*,*c*_ be a single cell dataset of features *f*_1_, *f*_2_,…, *f*_*n*_ by cells *c*_1_,*c*_2_,…,*c*_*m*_ and *Y*_*f*,*d*_ be a single cell dataset of the same features *f*_1_, *f*_2_,…, *f*_*n*_ by cells *d*_1_,*d*_2_,…,*d*_*p*_. Because the total number of cells that are measured in these experiments is generally much larger than the total number of features shared between the datasets, we opt for a diagonalized CCA implementation that has shown promising performance in related high-dimensional applications^70–72^. The goal is to find projection vectors *u* and *v* such that the correlation between the two indices *u*^*T*^ *X* and *v*^*T*^*Y* is maximized.

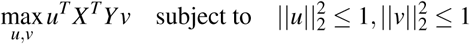

To find the canonical correlation vectors, we first standardize *X* and *Y* to have a mean of 0 and variance of 1. We use a standard singular value decomposition (SVD) to solve for the canonical correlation vectors *u* and *v* as follows:

Let

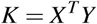

Decompose K via SVD:

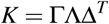

Where

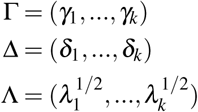

The canonical correlation vectors can then be obtained as the left and right singular values from the SVD for *i* = 1,…,*k*.

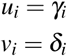

For computational efficiency, we approximate the SVD using the augmented implicitly restarted Lanczos bidiagonalization algorithm implemented in the irlba R package^73^. This allows us to obtain a user-defined number (*k*) of singular vectors that approximate the canonical correlation vectors (CCV). As described above, in this manuscript we set *k* to represent the “dimensionality” of the dataset.

Canonical correlation vectors (CCV) project the two datasets into a correlated low-dimensional space, but global differences in scale (for example, differences in normalization between datasets) can still preclude comparing CCV across datasets. To address this, we perform L2-normalization of the cell embeddings, where N is a vector of cell embeddings across the *k* CCV.

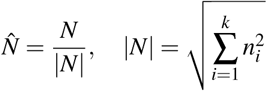

Following dimensional reduction, we identified the K-nearest neighbors (KNNs) for each cell within its paired dataset, based on the L2-normalized CCV. Finally, we identify mutual nearest neighbors (MNN; pairs of cells, with one from each dataset, that are contained within each other’s neighborhoods). We refer to these pairwise correspondences as “anchors", and wish to again highlight the foundational work of Haghverdi et al.^26^ for inspiring this concept. The size of this neighborhood (k.anchor parameter in FindTransferAnchors and FindIntegrationAnchors) was set to 5 for all analyses in this manuscript.

### Anchor scoring

The robust identification of anchor correspondences is key for effective downstream integration. Incorrect anchor pairs representing cells from distinct biological states can lead to incorrect downstream conclusions. In particular, cells that represent a biological state unique to one dataset should theoretically not participate in anchor pairs, yet in practice, they will do so with low frequency (Figure 1). Incorrectly identified anchors are similar to aberrant edges that can arise in KNN graphs (deemed ’short-circuits’ by Bendall et al.^30^) We therefore implement two steps (filtering and scoring anchors) to mitigate the effects of any incorrectly identified anchors.

First, we ensure that the anchors we identify in low-dimensional space also are supported by the underlying high-dimensional measurements. To do this, we return to the original data and examine the nearest neighbors of each anchor query cell in the reference dataset. We perform the search using the max.features (200) genes with the strongest association with previously identified CCV, using the TopDimFeatures function in Seurat, and search in L2-normalized expression space. If the anchor reference cell is found within the first k.filter (200) neighbors, then we retain this anchor. Otherwise, we remove this anchor from further analyses. We do not include a mutual neighborhood requirement for this step, as it is primarily intended as a check to ensure that we do not identify correspondences between reference and query cells with very divergent expression profiles. This procedure is uniformly applied with default parameters (max.features=200, k.filter=200), for all analyses in this manuscript.

Additionally, to further minimize the influence of incorrectly identified anchors, we implemented a method for scoring anchors that is similar to the use of shared nearest neighbor (SNN) graphs in graph-based clustering algorithms. By examining the consistency of edges between cells in the same local neighborhood, SNN metrics add an additional level of robustness to edge identification^31^. For each reference anchor cell, we determine its k.score (30) nearest within-dataset neighbors and its k.score nearest neighbors in the query dataset. This gives us four neighbor matrices that we combine to form an overall neighborhood graph. For each anchor correspondence, we compute the shared neighbor overlap between the anchor and query cells, and assign this value as the anchor score. To dampen the potential effect of outlier scores, we use the 0.01 and 0.90 quantiles to rescale anchor scores to a range of 0 to 1.

We find that when ground truth data is available for evaluating anchors, anchors representing reference and cell pairs have significantly higher scores than incorrect anchors (Figure 2I). Therefore, in downstream calculations (see below), anchors with lower scores are downweighted in favor of anchors with higher scores. The k.score parameter is fixed to 30 for all analyses in this manuscript. This procedure is implemented in the ScoreAnchors internal Seurat function, which is called by FindIntegrationAnchors or FindTransferAnchors in Seurat.

### Anchor weighting

We construct a weight matrix *W* that defines the strength of association between each query cell *c*, and each anchor *i* . These weights are based on two components: the distance between the query cell and the anchor, and the previously computed anchor score. In this way, query cells in distinct biological states (for example alpha cells and gamma cells) will be influenced by distinct sets of anchors, enabling context-specific batch correction. Additionally, robust anchors (with high scores) will gain influence across the query dataset, while inconsistent anchors will be downweighted. For each cell *c* in the query dataset, we identify the nearest *k.weight* anchors cells in the query dataset in PCA space. Nearest anchors are then weighted based on their distance to the cell *c* over the distance to the *k.weight*-th anchor cell and multiplied by the anchor score (*S*_*i*_). For each cell *c* and anchor *i*, we first compute the weighted distances as:

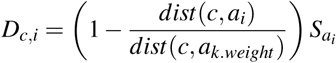

We then apply a Gaussian kernel:

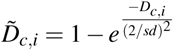

where *sd* is the Gaussian kernel bandwidth, set to 1 by default. Finally, we normalize across all *k.weight* anchors:

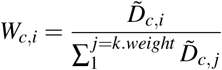

For identifying anchors for integration, we set k.weight = 100. For identifying transfer anchors, we set k.weight = 50. We reasoned that the batch vector information may be similar for closely related cell types, and so opt to take into account batch information for more anchors in integration analyses. In contrast, label information for different but closely related cell types would not improve the accuracy of cell type predictions, and so we consider a smaller number of anchors surrounding each cell. This procedure is implemented in the FindWeights internal Seurat function, which is called by IntegrateData or TransferData.

### Data integration for reference assembly

Once we have identified anchors and constructed the weights matrix, we follow the strategy outlined by Haghverdi et al.^26^ for batch correction. We first calculate the matrix *B*, where each column represents the difference between the two expression vectors for every pair of anchor cells,*a* :

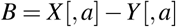

We then calculate a transformation matrix, *C*, using the previously computed weights matrix and the integration matrix as:

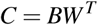

We then subtract the transformation matrix, *C*, from the original expression matrix,*Y*, to produce the integrated expression matrix *Ŷ*:

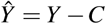

This step is implemented in the IntegrateData function in Seurat. The corrected expression matrix can be treated as a single normalized scRNA-seq matrix, and can be processed downstream using any single cell analytical toolkit. Notably, in Seurat, we continue to store the original uncorrected expression data, to facilitate downstream comparisons across datasets.

### Multiple Dataset Integration

Our approach to multiple dataset integration draws inspiration from methods for multiple sequence alignment. Many multiple sequence alignment algorithms begin with the construction of all pairwise alignments and proceed to merge these pairwise alignments to progressively form the final multiple sequence alignment^33^. Here, we use a similar approach where we first identify and score anchors between all pairs of datasets and then progressively build the final integrated dataset.

To integrate multiple datasets, we first determine the order in which to merge the datasets after pairwise anchor identification. To do this we first define a distance between any two datasets as the total number of cells in the smaller dataset divided by the total number of anchors between the two datasets. We compute all pairwise distances between datasets and then perform hierarchical clustering on this distance matrix using the hclust function from the stats R package. This returns a guide tree which we use to iteratively merge the datasets using the integration procedure described above to form the final integrated dataset. This procedure is implemented in the IntegrateData function in Seurat.

### Label Transfer

For cell metadata transfer, we create a binary classification matrix *L* containing the classification information for each anchor cell in the reference dataset. Specifically, each row in *L* corresponds to a possible class and each column corresponds to a reference anchor. If the reference cell in the anchor belongs to the corresponding class, that entry in the matrix is filled with a 1, otherwise the entry is assigned a 0. We then compute label predictions, *P*_*l*_, by multiplying the anchor classification matrix *L* with the transpose of the weights matrix *W*:

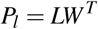

This returns a prediction score for each class for every cell in the query dataset that ranges from 0 to 1, and sums to 1.

### Feature Imputation

Our procedure for transferring continuous data is closely related to discrete label transfer. We compute new feature expression predictions, *P*_*f*_, by multiplying a matrix of anchor features to be transferred, *F*, with the transpose of the weights matrix *W*:

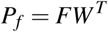

This returns a predicted expression matrix for each feature (row) in *F* for each cell in the query dataset. Feature imputation and label transfer are both implemented in the TransferData function in Seurat.

### Processing of single cell datasets

#### Data Acquisition and QC

The data used for the majority of the analyses in this paper come from publicly available repositories and data portals, and we are grateful to all the groups and organizations for making their data readily accessible. We obtained the human pancreatic islet datasets from the following accession numbers: GSE81076 (CelSeq), GSE85241 (CelSeq2), GSE86469 (Fluidigm C1), E-MTAB-5061 (SMART-Seq2), and GSE84133 (inDrops). We filtered out cells for which fewer than 1,750 unique genes/cell (Celseq) or 2,500 genes/cell (CelSeq2/Fluidigm C1/SMART-Seq2) were detected. For the inDrops data sets, we kept all cells with previously annotated cluster information. We obtained the UMI count matrix for the mouse retinal bipolar cell dataset under the accession number GSE81904, keeping only those cells with previously annotated cluster information. The *Tabula Muris* datasets were obtained from FigShare for the Version 1 release^74,75^. The human bone marrow dataset was obtained from the Human Cell Atlas Data Portal preview site^49^. We filtered out any cells for which fewer than 500 genes were detected and any genes that were expressed in fewer than 100 cells. The osmFISH data was obtained from the Linnarsson Lab website^76^. The mouse visual cortex SMART-seq4 data was obtained from the Allen Brain Data Portal^45,77^. Any cells that were annotated as either ’Low Quality’ or ’No Class’ were removed. The mouse prefrontal cortex scATAC-seq gene activity scores and ATAC peak matrices were obtained from the Seattle Organismal Molecular Atlases (SOMA) Data Portal^78^. The STARmap 1,020-gene datasets from the mouse visual cortex were downloaded from the original paper’s companion website^16,79^. We kept all cortical cells, based on the provided class labels, and did not perform additional filtration based on total RNA counts observed per cell.

### Bone marrow mononuclear cells CITE-seq experiment

Bone marrow mononuclear cells from a single human donor were purchased from AllCells (cat : ABM007F, lot :3008803, donor age: 25-30yo). The day of the experiment, cells were thawed according to manufacturer’s protocol. Briefly, cell vials were sprayed with ethanol and placed in a 37°C water bath for 2 minutes to thaw. RPMI 10% media was used to wash and resuspend cells. Cell numbers and viability were estimated using trypan blue. Cells were resuspended in CITE-seq^3^ staining buffer (2%BSA/0.01%Tween in PBS) and incubated with FcX blocking reagent for 10 minutes (BioLegend, cat #: 422302) to block nonspecific antibody binding. Following FcX blocking, cells were incubated with a pool of 25 antibodies (1*µ*g/antibody) for 30 minutes at 4°C. To ensure we can accurately identify cell doublets and distinguish empty droplets from cells with low gene counts, cells were split into 10 tubes each containing a unique hashing antibody from BioLegend^80^ and were incubated at 4°C for an additional 20 minutes. After incubation, cells were washed three times with 1 mL of staining buffer to remove any unbound antibodies. At the end of the final wash, cells were passed through a 40*µ* m filter to remove cell clumps (VWR, cat #: 10032-802) and resuspended in 1xPBS at the appropriate cell concentration for 10X genomics 3’ scRNA-seq^81^.

#### Antibody List

The following human Totalseq BioLegend antibodies were included in the pool: CD3 (cat #: 300475), CD56 (cat #: 362557), CD19 (cat #: 302259), CD11c (cat #: 371519), CD38(cat #: 102733), CD45RA, (cat #: 304157) CD123(cat #: 306037), CD127 (cat #: 351352), CD4 (cat #: 300563), CD8a (cat #: 301067), CD14(cat #: 301855), CD16(cat #: 302061), CD25 (cat #: 302643), CD45RO (cat #: 304255), CD69 (cat #: 310947), CD197 (cat #: 353247), CD161 (cat #: 339945), CD28 (custom made, clone: CD28.2), CD27 (cat #: 302847), HLA-DR (cat #: 307659), CD57 (custom made, clone: QA17A04), CD79b (cat #: 341415), CD11a (cat #: 350615), CD34 (cat #: 343537). For cellular hashing the following TotalSeq hashtag antibodies were purchased from BioLegend: Hashtag 1 (cat #: 394601), Hashtag 2 (cat #: 394603), Hashtag3 (cat #: 394605), Hashtag 4 (cat #: 394607), Hashtag 5 (cat #: 394609), Hashtag 6 (cat #: 394611), Hashtag 7 (cat #: 394613), Hashtag 8 (cat #: 394615), Hashtag 9 (cat #: 394617), Hashtag 10 (cat #: 394619). For the complete list of antibody barcode sequences see Tables S5 and S6.

#### CITE-seq data preprocessing

CITE-seq RNA reads were mapped to the human genome (GRCh38) and transcripts quantified using CellRanger v2.1.0^81,82^. Antibody counts for CITE-seq^3^ and cell hashing^80^ were counted using CITE-seq-count (https://github.com/Hoohm/CITE-seq-Count). Antibody-derived tags (ADTs) and hashtag oligos (HTOs) for each cell were normalized using a centered log ratio (CLR) transformation across cells, implemented in the function NormalizeData with method=”CLR”, across=”cells” in Seurat v3. Cells were demultiplexed using the HTOdemux function in Seurat, and cell doublets and background empty droplets subsequently removed. RNA counts for each cell were then preprocessed as described above (Data preprocessing).

#### CITE-seq cross-validation

We separated the 33,454-cell CITE-seq dataset into two equal groups at random to produce a query and reference dataset for cross-validation of protein expression transfer accuracy between experiments. Within the query dataset, we removed and stored the measured protein expression data for each cell. We then ran our transfer workflow on the query and reference dataset with default parameters, transferring the protein expression values from the reference dataset onto the query. We then computed, for each protein in each query cell, the Pearson correlation between the predicted protein expression and the measured expression level.

To assess the relationship between the number of RNA features (genes) used to identify anchors between the datasets and the resulting accuracy, we first ranked each gene in the CITE-seq dataset by it’s contribution to the overall variance in the dataset by multiplying the gene’s PCA loading with the variation explained by the component. We then took increasing subsets for these genes starting with the highest-ranked genes, ranging from 10 to 1,000 genes in steps of 10, and repeated the cross-validation.

#### Protein expression transfer to the HCA

To transfer cell surface protein expression data to the Human Cell Atlas dataset, we first computationally removed doublets from each scRNA-seq batch using Scrublet^83^, then integrated the eight human bone marrow datasets (eight different donors) using the Seurat v3 integration method (FindIntegrationAnchors and IntegrateData functions in Seurat v3) with default parameters, a dimensionality of 50, and an eps=1 as described above. We then transferred protein expression from the 33,454-cell CITE-seq dataset to the 274,599-cell HCA dataset using the FindTransferAnchors and TransferData functions in Seurat v3.

#### Analysis of CD69+ bone marrow population

We identified a population of predicted CD69+/CD8+ cells in the HCA bone marrow dataset. To identify a gene expression signature associated with this group of cells, we first performed an initial clustering of the data using Louvain clustering based on a shared nearest neighbor graph, as implemented in the FindClusters function in Seurat with default parameters. We then isolated the cluster that contained a mixture of CD69+ and CD69-CD8+ cells. We further subdivided this cluster into CD69-high and CD69-low cells by fitting a three-component mixture model using the predicted CD69 expression data, using the normalmixEM function in the mixtools R package^84^. After grouping the cells into high- and low-expressing CD69 populations, we searched for differentially expressed genes between the two populations using the original (uncorrected) HCA scRNA-seq data. We used the logistic regression differential expression test^85^ implemented in the FindMarkers function in Seurat, with the donor as a latent variable (latent.vars=”orig.ident”, test.use=”LR”). We retained the top 25 differentially expressed genes based on highest fold-change expression. We performed gene ontology enrichment analysis on this set of genes for both molecular function and biological process, using the R package GOstats with a p-value cutoff of 0.001^86^.

#### Validation of CD69+ T cell population

To validate the CD69+/CD8+ T-cell population identified through our integration method, we performed bulk RNA-seq experiments on FACS-sorted cell populations from the same bone marrow cell sample. Bone marrow mononuclear cells were thawed as described above. Cells were resuspended in MACS buffer (2%BSA/2mM EDTA in PBS) and incubated with FcX blocking reagent for 10 minutes (BioLegend, cat : 422302) to block nonspecific antibody binding. Cells were stained with the following FACS antibodies: FITC-CD3 (clone HIT3a, 300306), APC-CD4 (clone RPA-T4, cat : 300514), APCCy7-CD8 (clone RPA-T8, cat : 301015) and PE-CD69 (clone FN50, cat : 310905). DAPI was used to exclude dead cells (Thermo Fisher Scientific, cat : D1306). CD4-/CD8+/CD3+/CD69+ and CD4-/CD8+/CD3+/CD69-cells were sorted into tubes (4 replicates per population, 300-3000 cells per replicate) containing RLT lysis buffer (Giagen, cat : 79216) using the SONY SH800 sorter. To remove cellular debris, AmPure bead cleanup was performed on all sample lysates. Reverse transcription, cDNA amplification and RNA-seq libraries were prepared as described previously^87^. To identify differentially-expressed genes between the CD69+ and CD69-sorted populations, we used DESeq2^88^ and filtered for significant genes with a log_2_-fold change in expression greater than 1.5 and a q-value of less than 0.01^89^.

### Calculation of Moran’s I

To calculate Moran’s I statistic, a measure of spatial autocorrelation, we compute:

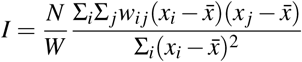

Where *N* is the number of spatial units (*i* and *j* for 2-dimensional space), *x* is the gene of interest, 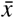 is the mean expression of gene *x*, *w*_*ij*_ is a spatial weight matrix with zeros on the diagonal, and *W* is the sum of all *w*_*ij*_. We computed the spatial weight matrix using the function dist in the R package stats, and applied a Gaussian kernel to the distance matrix to produce a smooth distribution. We used the implementation of Moran’s I available in the R package ape^90^, and acknowledge the Trapnell Lab Monocle 3 tutorials for suggesting the use of Moran’s I to estimate spatial autocorrelation in single cell data. We applied Moran’s I in two analyses: examining spatial (2D) patterns of gene expression in the mouse brain, and examining the dependence of gene expression on a 1D axis defined by CD69 expression in human bone marrow cells. In each case, we binned gene expression values for cells in small spatial regions before calculating spatial weights and Moran’s I.

### Assignment of cell type labels for pancreatic islet cells

To assign a set of consistent cell type labels to the pancreatic islet cell datasets, we based our classifications on the labels provided in the inDrops dataset. We first computed a PCA on the scaled integrated data matrix and used the first 30 PCs to build an SNN graph using the FindNeighbors function in Seurat with k.param set to 20. We then clustered the data using FindClusters in Seurat with the resolution parameter set to 1.5. For each resulting cluster, we assigned a label based on the most frequently occurring cell type in that cluster from the inDrops dataset (Table S2).

### Identification of rare subtypes in the *Tabula Muris* dataset

We integrated the two *Tabula Muris* datasets using the Seurat v3 integration method (FindAnchors and IntegrateData) with a chosen dimensionality of 100. We then normalized, scaled, and performed PCA on the integrated data as described in the Data preprocessing section above. The first 100 PCs were then used to construct an SNN matrix using the FindNeighbors function in Seurat v3 with k.param set to 20. We then identified clusters using the FindClusters command with the resolution parameter set to 4, identifying 132 total clusters. We annotated cluster 126 as mesothelial cells and cluster 124 as plasmacytoid dendritic cells based on the expression of known cell type markers (Figure S3E-F).

### Integration with simulated cell type holdouts

For both the pancreas and bipolar datasets, we performed a simulated holdout experiment where one cell type was completely removed from each dataset being integrated. These combinations are detailed in Table S1. After the cell type removal, highly variable genes were recalculated and integration features were selected. These features were then used as input to the integration procedure with the same default parameter settings as used in the full dataset integration.

We also tested the following existing integration methods on the same holdout datasets: Seurat v2^25^, mnnCorrect^26^, and scanorama^34^. For Seurat v2, we used the same feature set as determined for Seurat v3 to run a multi-CCA analysis followed by alignment (RunMultiCCA and AlignSubspace in Seurat v2). We used the first 30 aligned CCs to define the integrated subspace for clustering, visualization, and computing the integration metrics.

For mnnCorrect, we used the mnnCorrect function from the scran^91^ R package with the log-normalized data matrices as input, subset to include the same variable integration features we used for Seurat v3, and setting the pc.approx parameter to TRUE. This returned a corrected gene expression matrix on which we performed principle component analysis and kept the first 30 PCs as input for clustering, visualization, and computing the integration metrics.

For scanorama, we used the “correct” function with default parameter settings to batch correct the data and return an integrated expression matrix. The downstream processing here was kept the same as for mnnCorrect.

### Integration Metrics

To compare the results of the holdout integration experiments, we computed three measures of integration quality: the silhouette coefficient^92^, a mixing metric, and a local structure metric.

#### Silhouette coefficient

We computed the silhouette coefficient using the cluster package in R. Here, distances were computed in PCA space defined by the first 30 components for all methods except for Seurat v2, where we used the first 30 aligned CCs to define cluster distances. Clusters were defined using the previously assigned cell-type labels (Assignment of cell type labels for pancreatic islet cells). The silhouette coefficient gives a score for each cell that assesses the separation of cell types, with a high score suggesting that cells of the same cell type are close together and far from other cells of a different type. The silhouette score *s*(*i*) is defined for each cell *i* as follows. Let *a*(*i*) be the average distance of cell *i* to all other cells within *i* ’s cluster and *b*(*i*) be the average distance of *i* to all cells in the nearest cluster to which *i* does not belong. *s*(*i*) can then be computed as:

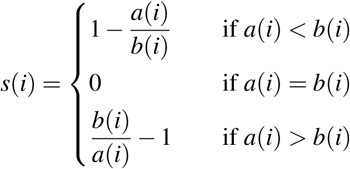

#### Mixing metric

We designed a “mixing metric” to evaluate how well mixed the input datasets were after integration. We first considered using a metric based on dataset entropy within an individual cell’s local neighborhood^93^. However, the assumptions of these methods are violated in cases where the distribution of cell type frequencies differs significantly across datasets, as is the case in many of our experiments. As an alternative, we reasoned that if the local neighborhood for a cell is well mixed, its closest neighbors should contain at least a small number (k = 5) of cells from each dataset. If the cell is poorly mixed, then its closest neighbors will likely stem only from a small subset of datasets (perhaps only its own). For each cell, we therefore examine the (k.max = 300) ranked nearest neighbors across all datasets. We also compute the k=5 closest neighbors for each dataset individually. We then ask which rank in the overall neighborhood list corresponds to the 5th neighbor in each dataset (with a max of 300), and took a median over all these values. This corresponds to a mixing metric per cell, and we averaged across all cells to obtain an overall mixing metric for each method. We found that this metric accurately reflected the the mixing of shared biological states across datasets, even when cluster frequencies differed. This metric is implemented in the MixingMetric function in Seurat v3.

#### Local structure metric

We computed a metric designed to determine how well the original structure of each dataset was preserved after integration. Here, we split the data back into its original datasets, re-compute an principal component analysis on the uncorrected data, and identify the k=100 closest nearest neighbors. We also computed the 100 nearest neighbors based on a principal component analysis of the integrated dataset. For every cell, we then looked at the intersection of these two neighborhoods and computed the fraction of overlap. For an overall score, we took the mean overlap fraction for all cells. This metric is implemented in the LocalStruct function in Seurat v3.

### Transferring cell type labels onto scRNA-seq data

In order to benchmark the projection of new data onto an existing reference, we performed the following experiments with the pancreas and bipolar datasets:

1. We generated 166 evaluation datasets for benchmark comparison. For each case, we removed one dataset from the reference to use as a query. We also removed all instances of one celltype from the reference (’withheld class’). We did this for all possible combinations of holdout datasets and cell types.
2. To ensure that a single cell type did not dominate in downstream evaluation, we downsampled the query dataset to contain a maximum of 100 cells per celltype. We then added or subtracted additional instances of cells in the “withheld” class, so that it composed 20% of the query.
3. We then integrated the reference using the same default workflow and parameter settings as for all previous integrations.

To classify cells using the Seurat v3 workflow, we first integrated the reference dataset using default parameters. We then classified query cells using the FindTransferAnchors and TransferData functions in Seurat with default parameters. We examined projection scores and assigned the cells with the lowest 20% of values to be “Unassigned".

We also repeated the classification using two functions from the scMAP R package: scmapCluster and scmapCell^42^. For these tests, we selected features using the selectFeatures function in scMap with n_features specified as 500. For scmapCluster, we set the similarity threshold parameter to -Inf to force assignments where possible. We then took the cells with the lowest 20% of similarity values and called them “Unassigned".

### scATAC-seq analysis

#### Preprocessing scATAC-seq data

We obtained scATAC-seq gene activity score and binarized peak count matrices for the mouse prefrontal cortex^18^. For all integration with scRNA-seq data, we used the gene activity score matrix. For finding differentially accessible peaks between groups of cells, we used the binarized peak count matrix. The authors instruct that scATAC-seq gene activity score matrix must be preprocessed and filtered, so we applied log-CPM (counts-per-million) normalization, and removed cells with less than 5,000 total peaks detected in the binary peak matrix.

#### Transferring cell type labels onto scATAC-seq cells

We found anchors between the pre-processed scATAC-seq cells (gene activity matrix) for the mouse prefrontal cortex and scRNA-seq cells from the mouse visual cortex^44,45^. We first found highly variable features in the scRNA-seq data using the FindVariableFeatures function in Seurat v3, as described above (Data preprocessing). We used the top 5,000 variable features that were also present in the scATAC-seq data as input to the integration, resulting in ~3,000 variable features as recommended by the original authors for downstream analysis^18^. We found anchors between the two datasets using the FindTransferAnchors function in Seurat v3, with the parameters dims=1:20 and reduction=”cca”. We transferred cell type labels from the scRNA-seq dataset to the scATAC-seq cells using the TransferData function in Seurat v3, with the parameters dims=1:20, reduction=”cca”.

To visualize the two datasets together, we transferred scRNA-seq data onto the scATAC-seq cells, using the same anchors as previously identified. We accomplished this by applying the same procedure used to impute transcriptome-wide expression in the STARmap dataset (see below). After imputation, we concatenated this matrix with the scRNA-seq dataset, performed a single PCA on both datasets, projected to two dimensions with UMAP, and colored the cells by their classification label. We emphasize that step is intended only for visual interpretation, and it is not necessary for us to jointly visualize the datasets, or transfer scRNA-seq data, in order to classify the scATAC-seq cells.

#### Identification of differentially accessible peaks and overrepresented motifs

We identified differentially accessible peaks between groups of scATAC-seq cells simply by ordering peaks by their fold-change accessibility between the groups, and retaining the top 1,000 peaks that displayed the greatest fold-change in accessibility. We searched for overrepresented DNA sequence motifs in accessible regions using the Homer package^94^, using the findMotifsGenome.pl program with default parameters, and the mm9 genome.

#### Psuedo-bulk ATAC-seq data collation

We split the scATAC-seq binary sequence/alignment map (BAM) file for the prefrontal cortex into individual files for each predicted cell type to create pseudo-bulk ATAC seq datasets for each celltype. We used custom Python code using the pysam package (https://github.com/pysam-developers/pysam) to extract scATAC-seq reads by their cell barcode^95^. We created normalized read coverage tracks (bigwig format) for each BAM file using the program bamCoverage in the deepTools package^96^ with the binSize parameter set to 5 and using the reads per kilobase per million mapped reads (RPKM) normalization option.

### Projecting gene expression and cell type labels onto spatially-resolved cells

#### Preprocessing STARmap data

We obtained STARmap gene count matrices and cell position information for two combinatorially-encoded 1,020-gene experiments from the STARmap companion website (https://www.starmapresources.com/data/^16^), and preprocessed the gene expression matrices as described above (Data preprocessing), with a normalization scaling factor equal to the median RNA counts per cell. To visualize spatial patterns of gene expression, we identified cell locations and morphologies using Python code provided by the original authors (https://github.com/weallen/STARmap^16^). Before transferring transcriptome-wide gene expression data from the SMART-seq4 dataset^44,45^ to the STARmap cells, we first integrated the two STARmap replicates using the Seurat v3 integration method. First, we identified anchors between the STARmap datasets using the FindIntegrationAnchors function in Seurat v3, using all 1,020 genes as input to the CCA. We then integrated the datasets using the IntegrateData function.

#### Data transfer

We then transferred transcriptome-wide gene expression data from the SMART-seq4 dataset to the integrated STARmap datasets using the FindTransferAnchors (reduction=”cca”) and TransferData functions in Seurat v3. We ran TransferData twice, once to transfer transcriptome-wide gene expression measurements, and again to transfer cell type labels from the SMART-seq4 dataset, using the same set of anchors. For all genes shown in Figure 5B (*Cux2*, *Lamp5*, *Rorb*, *Rab3c*, *Syt6*, *Sox2ot*, *Bsg* and *Sst*), imputations represent leave-one-out cross-validation of the STARmap data transfer. Specifically, we performed feature transfer independently for each gene, each time removing the gene of interest from the set of genes used to identify anchor cells between the STARmap and SMART-seq4 datasets.

#### Analysis of integrated STARmap data

After predicting cell type labels in the STARmap cells by transferring labels from the SMART-seq4 dataset, we observed 1,915 cells with high-confidence cell type predictions (prediction score > 0.5). To be more conservative, we chose to filter the predicted celltype labels to retain only the highest 50% scoring cells for each transferred cell type, retaining a total of 1,210 classified cells. We then performed differential expressed to identify gene expression markers that were upregulated in each classified cell type. We used a logistic regression test for differential expression^85^ on the uncorrected data with replicate as a latent variable, implemented in the FindMarkers function in Seurat (method=”LR”, latent.vars=”orig.ident”, assay=”RNA”).

To assess the robustness of our data transfer method, we first re-computed variable genes in each STARmap dataset, using the predicted gene expression data. Here, we selected the top 3,000 highly variable genes based on mean-variance dispersion. We did not use the variance-stabilizing transformation described above (Data preprocessing), as the predicted expression data are not discrete counts. For each gene identified as highly variable in either STARmap replicate, we calculated Moran’s I (see Calculation of Moran’s I, above), to estimate the relationship between predicted gene expression and spatial distribution for each gene. We then compared the Moran’s I value for each gene in the two replicates by calculating the Pearson correlation between Moran’s I values.

## Acknowledgements

This work was supported by an NIH New Innovator Award (1DP2HG009623-01; R.S.) and R01 (5R01MH071679-12; R.S.), Chan Zuckerberg Awards HCA-A-1704-01895 (R.S. and P.S.) and HCA2-A-1708-02755 (R.S) and an NSF Graduate Fellowship (DGE1342536; A.B.). We acknowledge members of the Satija Lab, NYGC Technology Innovation lab, Claude Desplan, Dan Littman, and Gord Fishell for helpful comments and discussion, and Josh Batson and Will Allen for sharing data and metadata.

## Author contributions

TS, AB, and RS conceived the research. TS and AB led computational work, assisted by PH, CH, and supervised by RS. EP and MS led experimental work, with assistance from WMM and supervised by PS. All authors participated in interpretation and writing the manuscript.

